# Disease dynamics during ecosystem restoration and faunal reassembly

**DOI:** 10.64898/2026.11.05.764583

**Authors:** Marina Voinson, Diana Erazo, Christina Faust, Nils Bunnefeld, Alfan A. Rija, Herwig Leirs, Luisa Páez-Triana, Vincent Sluydts, Joachim Marien, Lucinda Kirkpatrick

**Affiliations:** EVECO group, Univ. of Antwerp, Antwerp, Belgium; African Museum, Brussels, Belgium; School of Biodiversity, One Health and Veterinary Medicine, University of Glasgow, Glasgow, UK; School of Life and Biological Sciences, University of Stirling, Stirling, UK; Department of Wildlife Management, Sokoine University of Agriculture, Morogoro, Tanzania; Department of Biomedical sciences, Institute of Tropical Medicine Antwerp, Antwerp, Belgium; School of Environmental and Natural Sciences, Bangor University, Bangor, UK

## Abstract

Biodiversity recovery is increasingly promoted as a strategy for restoring ecosystem services, including the regulation of infectious diseases. However, it remains unclear how the potential for pathogen transmission interacts with the timing of biodiversity recovery and the life history of the recovering species to shape the disease dynamics during restoration. Here, we developed a stochastic multi-host metacommunity epidemic model to investigate how the timing of the establishment of slow life history species, pathogen transmissibility, and host-community succession jointly influence disease outcomes in recovering ecosystems. Pathogen transmissibility (*R*_0_) was the dominant predictor of epidemic intensity, whereas the delayed timing of establishment of slow life history species during restoration emerged as a major ecological determinant of pathogen persistence. Delayed establishment of slow life-history hosts consistently increased disease prevalence and persistence by prolonging the dominance of highly competent fast life-history hosts. Generalized additive models revealed a strong interaction between delayed timing of slow life-history species establishment and pathogen transmissibility, explaining more than 90% of the variation in pathogen persistence. The influence of timing of establishment of slow life history species was weak for pathogens with low transmission potential but increased markedly as *R*_0_ increased. Sensitivity analyses further showed that, once pathogen transmissibility was controlled, timing of establishment became the strongest predictor of persistence, whereas among-host spillover transmission primarily influenced the speed of epidemic spread. In contrast, maximum species richness had comparatively little influence on disease outcomes, indicating that the functional composition and assembly timing of recovering communities may be more important than richness alone for predicting disease dynamics. Together, these results demonstrate that disease regulation during ecosystem restoration depends not only on pathogen transmissibility, but also on the life history traits and the timing of establishment of species within recovering communities. Our findings identify life history traits and the timing of when functional communities are restored as a key ecological lever influencing long-term pathogen maintenance in restored ecosystems.

## 1 Introduction

Human-driven environmental change is reshaping ecological communities worldwide, accelerating biodiversity loss and altering the ecosystem processes that regulate infectious diseases [1, 2]. Habitat degradation and land-use change often favour fast-lived, generalist species that exhibit high pathogen competence, while disproportionately eliminating long-lived and specialised taxa that can contribute to disease regulation through dilution, encounter reduction, or community stabilisation mechanisms [3]. Consequently, biodiversity loss is increasingly recognised not only as a conservation crisis, but also as a potential driver of emerging infectious diseases and zoonotic spillover risk.

In response, ecosystem restoration has become a central pillar of global sustainability agendas. Initiatives such as the United Nations Decade on Ecosystem Restoration, the Bonn Challenge, and the European Nature Restoration Law aim to reverse biodiversity decline and recover ecosystem functioning at unprecedented scales [4, 5, 6]. Beyond benefits for biodiversity conservation, climate mitigation and ecosystem resilience, restoration has also been proposed as a nature-based solution for reducing infectious disease risk [7, 8]. Yet empirical evidence remains scarce. A recent synthesis identified only 39 studies explicitly examining links between ecological restoration and zoonotic disease risk, highlighting substantial uncertainty regarding how biodiversity recovery influences pathogen transmission [9, 10].

A major challenge is that restoration is inherently dynamic. Rather than representing a simple return to historical conditions, ecosystem recovery unfolds through trajectories of colonisation, succession, and community assembly [11, 12, 13]. Early successional communities are typically dominated by fast-living, disturbance-tolerant species characterised by high demographic turnover and broad ecological niches, whereas slower-living and more specialised species establish later as ecosystems mature [14, 15]. Because host life-history strategies influence pathogen transmission, competence and persistence [16, 17], disease risk is likely to vary throughout the restoration process. Consequently, recovering ecosystems may pass through transient phases of elevated pathogen amplification before disease regulation is restored.

Despite growing recognition of these dynamics, mechanistic frameworks linking restoration trajectories, community assembly and infectious disease processes remain limited [18]. Most studies evaluate the relationships between biodiversity and disease using static representations of ecological communities, overlooking the temporal processes through which biodiversity is restored [19, 20]. At the same time, pathogens differ markedly in their transmission potential, which determines their ability to invade and persist within host communities [21]. Although variation in pathogen transmissibility is known to influence epidemic dynamics, it remains unclear whether the effectiveness of restoration in mitigating disease risk depends on pathogen transmission potential [22]. Understanding how faunal reassembly during restoration interacts with pathogen transmissibility is therefore essential for predicting disease outcomes in recovering ecosystems.

Life-history theory provides a useful framework for understanding how community succession may influence infectious disease dynamics. Species occupying the fast end of the fast–slow continuum are typically characterized by rapid reproduction, short generation times, and high population turnover, whereas species at the slow end generally exhibit longer lifespans, lower reproductive rates, and greater investment in survival [23, 17]. These life-history differences are often associated with variation in pathogen competence and transmission. Fast life-history hosts frequently allocate proportionally fewer resources to long-term immune defence and can support higher pathogen replication or transmission rates, making them potentially important reservoirs for infectious diseases [24, 17, 16]. In contrast, slow life-history species are often associated with lower transmission potential and greater demographic stability. Because ecological restoration typically drives a gradual replacement of fast by slow life-history species during succession [14, 15], changes in host life-history composition may simultaneously reflect improving ecosystem condition and declining opportunities for pathogen amplification. Consequently, restoration trajectories may influence disease risk not only through changes in biodiversity, but also through shifts in the functional characteristics of host communities.

Here, we address this knowledge gap using a mechanistic multi-host epidemiological model that explicitly couples community assembly and disease dynamics during ecological restoration. We developed a stochastic metapopulation Susceptible–Infected–Recovered (SIR) framework that incorporates fast and slow-living host species, demographic turnover, interspecific transmission, and progressive recolonisation during ecosystem recovery. By combining global sensitivity analyses, machine-learning approaches, and mechanistic modelling, we assess how restoration timing, biodiversity recovery and pathogen life-history traits jointly shape epidemic outcomes. Specifically, we test three hypotheses. First, we hypothesise that restoration timing is a major determinant of disease dynamics because it controls when slow life-history hosts can exert regulatory effects on pathogen transmission. Second, we hypothesise that pathogen transmission potential (*R*_0_) is a primary determinant of epidemic intensity and persistence. Third, we hypothesise that restoration timing and pathogen transmissibility interact, such that the epidemiological consequences of community succession depend on pathogen transmission potential.

By identifying when and under which ecological conditions restoration reduces disease risk, this study provides a mechanistic basis for integrating infectious disease regulation into ecosystem restoration planning. More broadly, it contributes to emerging efforts to link biodiversity recovery, ecosystem functioning and health within a One Health framework.

## 2 Materials and Methods

### Overview of the modelling framework

We developed a stochastic, spatially implicit community model to examine how ecological restoration trajectories influence infectious-disease dynamics. Restoration changes host-community composition through sequential colonization, while host populations undergo demographic and epidemiological processes. Each established species is represented by a well-mixed population with susceptible (*S_i_*), infectious (*I_i_*), and recovered (*R_i_*) individuals, where *N_i_* = *S_i_* + *I_i_* + *R_i_*. Populations experience births, deaths, recovery, within-population transmission, and transmission between populations. The model distinguishes two stylized host strategies: early-successional, fast-life-history populations (denoted *r*) and later-establishing, slow-life-history populations (denoted *K*). Simulations were implemented in C++ using a fixed-step tau-leaping (*τ* = 0.01) approximation to the underlying continuous-time Markov process. The complete source code and simulation framework have been archived on Zenodo [25]. Full transition rules, restoration assumptions, and numerical implementation are provided in the Supplementary Methods.

### Restoration trajectory and ecological succession

The richness of the target species increases along a saturating restoration trajectory controlled by the colonization rate and the maximum richness *S*_max_. At each tau-leap update, the target richness is evaluated at the current simulation time; any difference between target and established richness is filled by sequential species introductions. Consequently, multiple species may be introduced at the same update when target richness rises by more than one species. These introductions are scheduled by the richness trajectory and are not stochastic birth or infection events. We represent ecological restoration as a gradual shift from communities dominated by early-successional, fast-life-history hosts (*r*) toward communities increasingly occupied by late-successional, slow-life-history hosts (*K*). The key restoration parameter is the introduction threshold *T_K_*, which determines when the establishment of *K* populations becomes possible. Ecologically, *T_K_* controls the duration of the early-successional phase and therefore how long the host community remains dominated by *r* populations before later-successional hosts begin to establish. Before *T_K_*, the probability of establishing a *K* population is zero and all newly introduced species belong to the *r* group. At and after *T_K_*, each arriving species is assigned to group *K* with probability

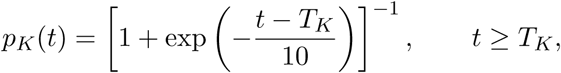

and to group *r* with probability 1 − *p_K_*(*t*). The scale parameter of 10 time units controls the speed of the transition. At the threshold, *p_K_*(*T_K_*) = 0.5, and the probability approaches one as succession proceeds. Consequently, earlier values of *T_K_* lead to earlier establishment of late-successional hosts and a faster compositional transition, whereas later values of *T_K_* prolong the dominance of *r* populations. When several species are added at the same richness update, their group identities are assigned sequentially using the current value of *p_K_*(*t*). This stochastic assignment reflects a gradual change in establishment opportunities during restoration rather than a deterministic replacement of all *r* populations by *K* populations. The first successful establishment of a *K* population triggers a one-time competitive transition intended to represent a major ecological shift associated with late-successional restoration. At this event, previously established *r* populations are reduced in abundance according to a competition parameter (*comp*), which determines the proportion of individuals removed from each compartment (*S*, *I*, and *R*). Subsequent *r* populations are initialized at a reduced abundance equal to (1− *comp*)*N_i,r_*, whereas newly arriving *K* populations are initialized at *N_i,K_*. Populations established before the transition retain their post-reduction abundances. These rules represent an imposed successional shift rather than an emergent model of resource competition. Because the magnitude of competitive replacement between fast and slow life-history hosts is uncertain, we performed an additional robustness analysis in which *comp* varied between 0.35 and 0.65 while all other assumptions were held constant (Fig. S5). This analysis did not qualitatively alter predictor rankings or the main epidemiological patterns, indicating that the principal conclusions were robust to uncertainty in the strength of competitive replacement. The first established host population receives the initial infection according to the initialization rule implemented in the simulation code. Overall, the restoration hypothesis represented by the model is that communities initially consist exclusively of fast-life history hosts, remain dominated by these hosts until the threshold *T_K_* is reached, and then progressively transition toward greater representation of slow-life history hosts. Earlier values of *T_K_* therefore correspond to earlier restoration-induced community turnover, whereas later values prolong the period during which epidemiologically competent *r* hosts dominate the system. Examples of community assembly trajectories and their associated epidemiological dynamics under early and delayed establishment of slow life-history hosts are shown in Fig. S1.

### Host strategies, demography, and transmission

The *r* and *K* labels denote functional abstractions along a fast–slow life-history continuum rather than discrete taxonomic or universal ecological categories. In the model, life-history differences are represented through group-specific demographic and epidemiological parameters. Fast life-history hosts (*r*) are characterised by higher demographic turnover and may differ from slow life-history hosts (*K*) in their initial abundance, birth and death rates, transmission coefficients, and recovery rates. These parameter differences are intended to capture generic contrasts between rapidly reproducing, disturbance-tolerant hosts and more slowly reproducing, longer-lived hosts. Within-population infection incidence is mass-action, depending on the product of susceptible and infectious abundance and the strategy-specific transmission coefficient. Between-population transmission is represented by infectious sources, normalized by the abundance of the relevant source group as detailed in the Supplementary Methods.

### Tau-leap simulations and epidemiological outcomes

At each fixed interval *τ*, transition counts are drawn from binomial or Poisson distributions using rates evaluated at the beginning of the interval, and the resulting changes are then applied to the populations. Competing transitions out of a compartment are sampled jointly to prevent removals from exceeding the number of individuals present at the start of the step. The restoration trajectory and strategy-assignment probability are updated at the start of each step. Tau-leaping is an approximation to the continuous-time stochastic process; its numerical adequacy is assessed by comparing outcomes across decreasing step sizes before final inference. For each realization, we calculate community-level prevalence as 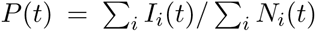 when total abundance is positive. We record peak prevalence, final prevalence, time to peak, whether infection was ever observed, and whether infection persists or is absent at the simulation end. A realization in which infection was never observed is distinguished from one in which infection occurred and subsequently became extinct. Demographic and restoration dynamics continue to the prescribed simulation horizon after infection extinction, so final community composition is evaluated at that horizon.

### Global sensitivity analysis

We used Latin hypercube sampling (LHS) to explore variation in epidemiological and restoration parameters. These included the basic reproduction number of pathogens circulating within fast life-history hosts (*R*_0_), maximum species richness (*S*_max_), the strength of competitive replacement associated with slow life-history host establishment, within-host transmission parameters (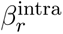 and 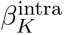), among-host transmission (*β*^inter^), recovery rate of slow life-history hosts (*γ_K_*), species establishment rate, and the timing of slow life-history host (*T_K_*). Unless otherwise stated, transmission and recovery parameters and species establishment rate were sampled over 0.5–1.5 times their reference values, whereas *T_K_* was sampled between 1 and 100 model-time units. In analyses designed to isolate ecological effects from variation in pathogen transmissibility, *R*_0_ was held constant and recovery rates were calculated from transmission parameters to satisfy the specified value of *R*_0_. Each sampled parameter set was simulated using a reproducible random seed derived from the parameter-set and replicate identifiers. The global sensitivity analysis was based on 1000 Latin hypercube parameter sets, each simulated with 100 stochastic replicates. For predictive sensitivity analyses, parameter-set-level separation was used for training and testing to prevent replicates from the same sampled parameter combination appearing in both datasets. Predictor rankings and validation metrics are therefore interpreted as measures of predictive importance within the sampled parameter space rather than as causal effects. Variable importance was quantified using XGBoost gain metrics, and robustness was evaluated using SHAP analyses and bootstrap resampling.

### Model assumptions and scope

The model assumes well-mixed populations, homogeneous parameters within each population, lifelong immunity following recovery (i.e. recovered individuals do not return to the susceptible class), and two stylized life-history strategies. Restoration is represented by a saturating richness trajectory and a probabilistic shift in the strategy of arriving populations after *T_K_*. The baseline model assumes that the first establishment of slow life-history hosts reduces the abundance of previously established fast life-history hosts, representing a simplified successional transition. Because the magnitude of this effect is uncertain, we additionally varied the strength of this reduction in a separate sensitivity analysis to assess the robustness of model predictions to assumptions about competitive replacement. This mechanism is intended to represent a major ecological shift associated with succession rather than a mechanistic model of resource competition. The model does not explicitly represent spatial dispersal, habitat quality, behavioural responses, pathogen evolution, waning immunity, or direct human exposure. There-fore, its inferences concern how restoration trajectories, host-community composition, and pathogen characteristics influence amplification and persistence of community-level infections. Figure 1 summarises the conceptual structure of the model, including faunal succession during restoration and the epidemiological dynamics associated with fast and slow life-history hosts.

**Figure 1:**
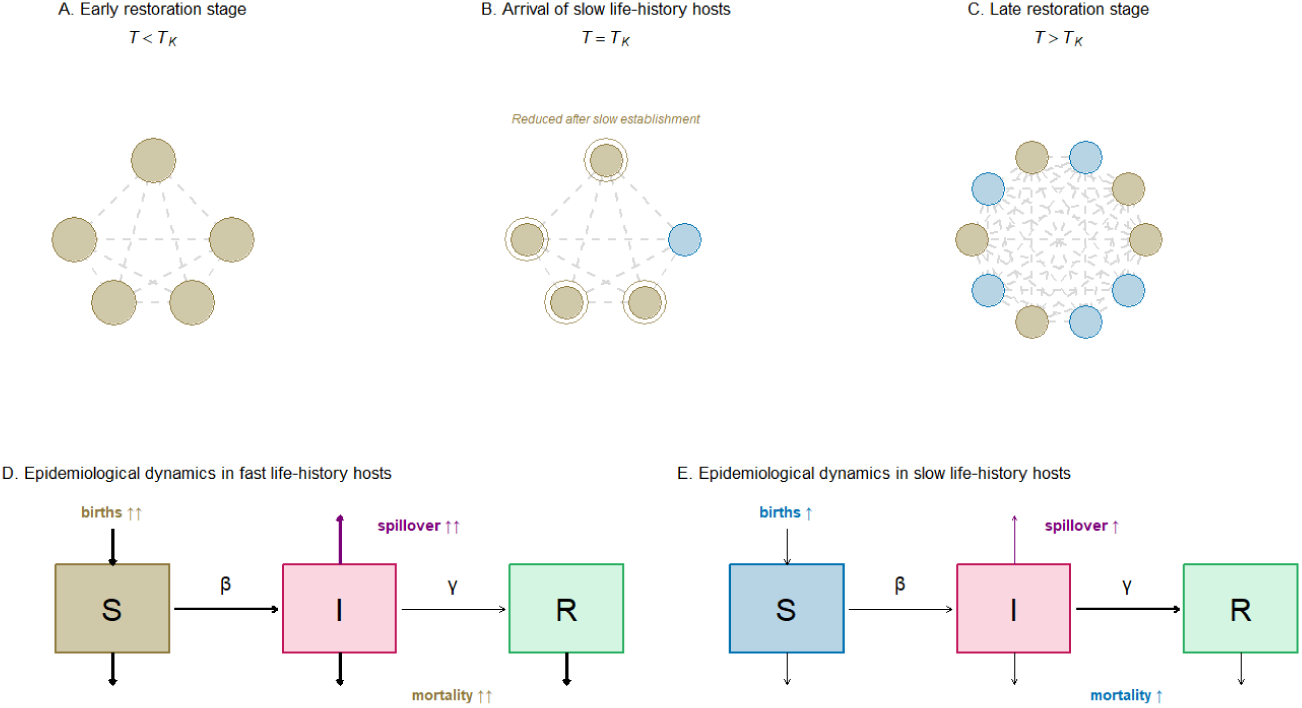
Conceptual framework linking faunal succession and disease dynamics during ecosystem restoration. (A–C) Community assembly during restoration. Early restoration stages are initially dominated by fast life-history host species (A). At the restoration threshold (*T_K_*), slow life-history hosts progressively establish, while existing fast life-history populations are reduced in size (B). Over time, communities transition toward a mixed assemblage containing both fast and slow life-history hosts (C). Dashed lines represent potential transmission pathways among host populations. (D–E) Schematic representation of epidemiological processes within fast and slow life-history host populations. Both host types follow susceptible–infected–recovered (SIR) dynamics, but differ in demographic and epidemiological characteristics. Fast life-history hosts are characterized by higher birth rates, stronger among-host transmission (spillover), and higher mortality, whereas slow life-history hosts exhibit lower demographic turnover and weaker spillover transmission. Ecological restoration therefore modifies disease dynamics by altering the relative contribution of fast and slow life-history hosts to pathogen transmission through time.

## 3 Results

### Drivers of disease dynamics

XGBoost analyses (Fig. 2) identified pathogen transmissibility in fast life-history hosts (*R*_0_ in (r species)) as the dominant predictor of all epidemiological outcomes. The importance of *R*_0_ was particularly strong for peak prevalence (89.8%), but remained high for persistence (66.5%), final prevalence (66.4%), and time to peak (40.9%). The timing of slow life-history hosts establishment (*T_K_*) played a secondary but consistent role. Its influence was strongest for pathogen persistence (25.1%) and weaker for final prevalence (11.3%) and time to peak (11.2%), while contributing little to variation in peak prevalence (1.0%). Transmission among fast life-history host populations (r species) had contrasting effects across epidemiological outcomes. It was relatively important for final prevalence (14.1%) and time to peak (23.2%), but had little influence on persistence (2.6%) or peak prevalence (1.4%). Maximum species richness explained only a small proportion of variation in final prevalence (3.7%) and persistence (1.3%), although its contribution was somewhat greater for time to peak (11.0%). Overall, richness effects were consistently weaker than those associated with pathogen transmissibility or restoration timing.

**Figure 2:**
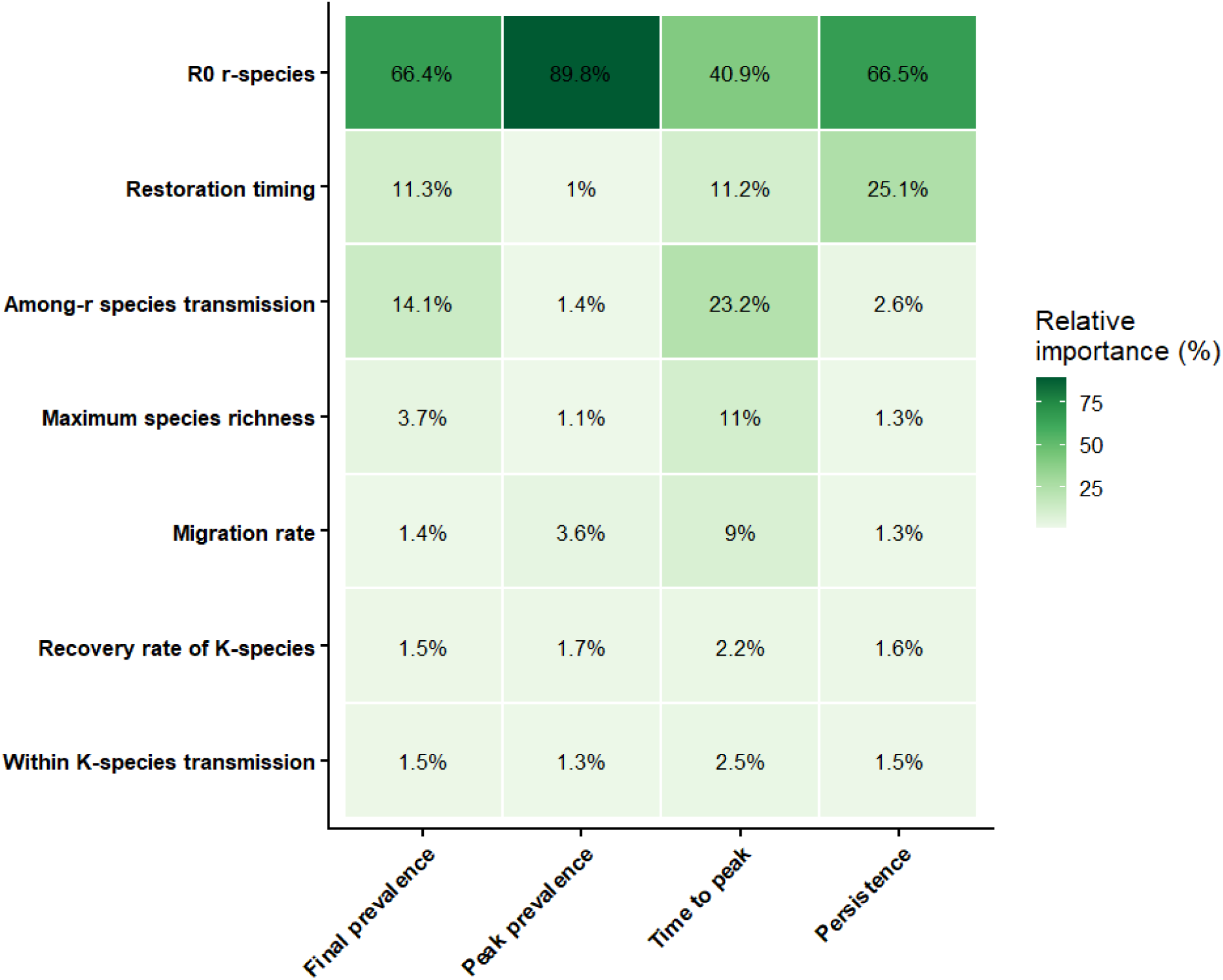
Relative importance of ecological and epidemiological predictors across four epidemiological outcomes, estimated using XGBoost. Cell values and colours indicate the relative contribution (%) of each predictor to model performance for final prevalence, peak prevalence, time to epidemic peak and persistence probability.

Parameters associated with slow life-history hosts (K species) contributed relatively little across all epidemiological outcomes (1.3–2.5%), indicating that disease dynamics were primarily governed by pathogen transmissibility in fast life-history hosts and by the timing of ecological succession.

### Interactive effects of restoration timing and pathogen transmissibility on persistence

Generalized additive models revealed a strong interaction between the timing of K-species establishment and pathogen transmissibility in determining pathogen persistence (Fig. 3). The interaction between the timing of K-species introduction (*T_K_*) and the basic reproduction number of pathogens circulating within fast life-history hosts (*R*_0_) was highly significant (*F* = 730.5, *p <* 0.001) and explained most of the variation in persistence probability (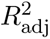 = 0.925; deviance explained = 92.6%). Persistence probability increased with both pathogen transmissibility and delayed establishment of K-species hosts. When pathogen transmissibility was low (*R*_0_ ≈ 2), persistence remained low regardless of the timing of K-strategist arrival and rarely exceeded 0.1. In contrast, when transmissibility was high (*R*_0_ ≈ 5), persistence exceeded 0.5 when K-species arrived later. Thus, the influence of restoration timing depended strongly on pathogen transmissibility. SHAP dependence analyses supported the same pattern (Fig. S4), indicating that the effect of delayed K-species introduction on pathogen persistence became progressively stronger as *R*_0_ increased.

**Figure 3:**
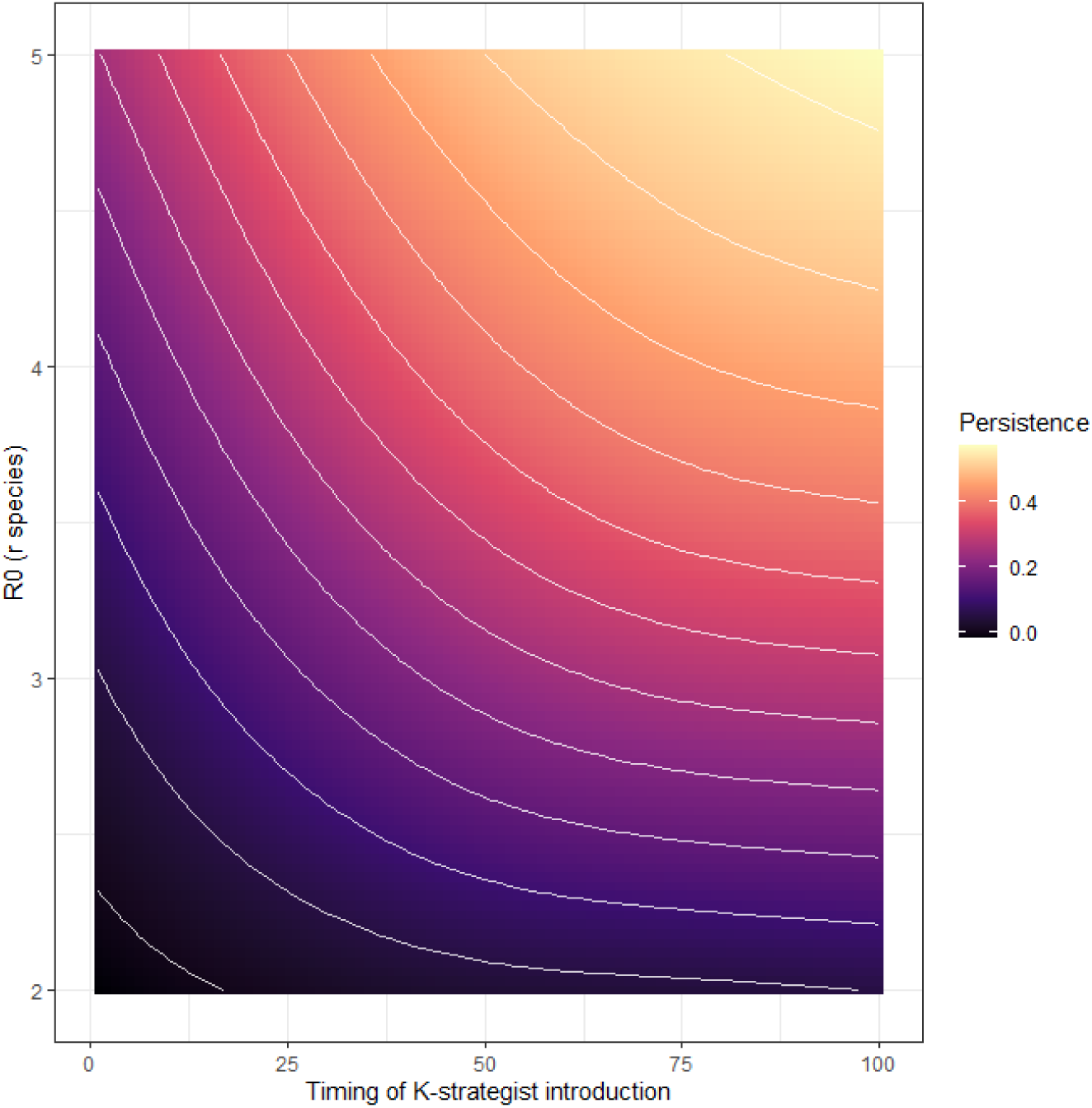
Interaction between restoration timing and pathogen transmissibility (R0) in determining pathogen persistence. The surface shows generalized additive model predictions across the explored parameter space. Colours indicate predicted persistence probability and contour lines denote equal predicted values.

To separate assemblage timing effects from variation in pathogen transmissibility, we repeated the analysis under a constant-*R*_0_ scenario (Fig. S3). The timing of establishment of K-species remained a major predictor of persistence, with its contribution increasing from 25.1% under variable *R*_0_ to 55.6% when *R*_0_ was held constant. This result indicates that ecosystem recovery influences pathogen persistence through changes in community succession and host composition rather than through variation in pathogen transmissibility alone.

Additional analyses indicated that the restoration effect could not be explained solely by variation in final community abundance or final community composition. When persistence probability was modelled as a function of K-species recovery together with final abundance, fast-host richness, or slow-host richness, the timing of K-species establishment remained highly significant whereas the community metrics themselves did not (see Tab S4). This suggests that pathogen persistence depends less on the final state of the community than on the sequence of ecological transitions leading to that state. In particular, the timing of K-species establishment appears to influence persistence by altering the duration for which competent fast life-history hosts dominate recovering communities, rather than through simple changes in abundance or richness.

### Spillover transmission delays epidemic spread

Generalized additive models showed that among-*r* -species transmission and the timing of K-species establishment jointly influenced the timing of epidemic spread across the community (Fig. 4). The interaction between among-*r* -species transmission (*β*_inter_) and the timing of K-species establishment (*T_K_*) was highly significant (*F* = 33.41, *p <* 0.001), although it explained a smaller proportion of variation than the persistence models (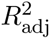 = 0.218; deviance explained = 22.4%). Time to epidemic peak increased with among-*r* -species transmission. At low values of *β*_inter_, epidemic peaks occurred relatively rapidly following pathogen introduction, whereas higher transmission among reservoir populations produced substantially later community-wide peaks. This pattern suggests that spillover among competent host populations sustains infection across multiple interconnected populations, extending epidemic growth rather than generating a rapid, synchronized outbreak. Delayed establishment of slow life-history hosts further increased time to peak by prolonging the period during which competent fast life-history hosts dominated the community. Additional analyses indicated that the effect of the timing of K-species establishment on epidemic timing was partially associated with final community structure (Table S4). In contrast to persistence, both final abundance and final host composition significantly contributed to variation in time to peak. Together, these results suggest that spillover primarily governs the temporal progression of epidemics, whereas the timing of K-species establishment modulates the duration of transmission opportunities through its effects on community assembly and the resulting host-community structure.

**Figure 4:**
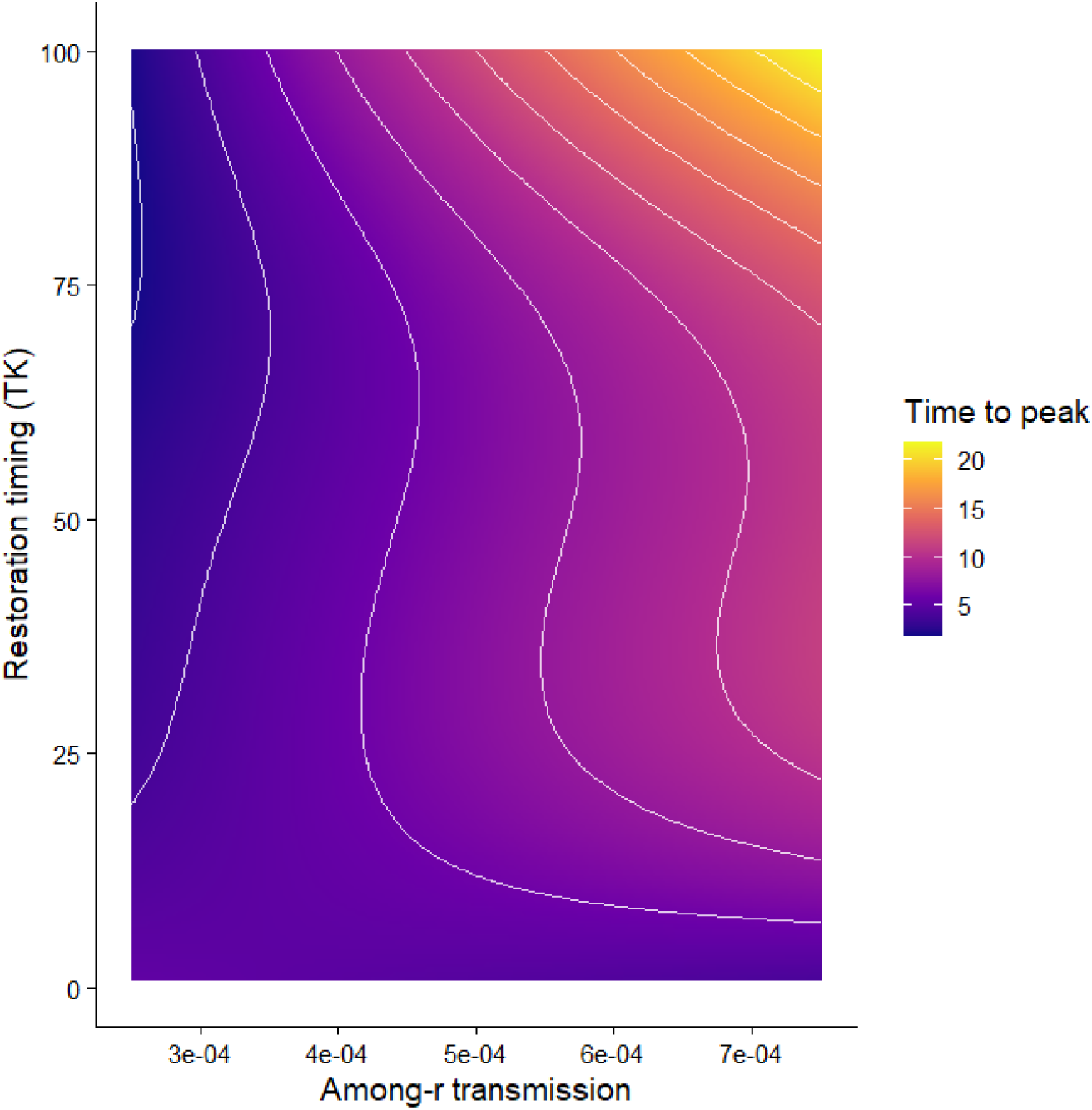
Interaction between among-*r* -species transmission (*β*_inter_) and restoration timing (*T_K_*) on the time to epidemic peak. Colours and contour lines show GAM-predicted values.

## 4 Discussion

Our results show that disease dynamics during faunal reassembly following ecosystem restoration are jointly determined by pathogen transmission potential and host-community succession. Across the explored parameter space, pathogen transmissibility (*R*_0_) emerged as the dominant predictor of epidemic intensity and prevalence, whereas the timing of slow life-history host establishment strongly influenced pathogen persistence. These findings indicate that disease outcomes cannot be predicted from pathogen traits or faunal reassembly trajectories independently. Instead, epidemiological dynamics emerge from the interaction between pathogen transmission potential and the temporal sequence of community assembly, extending previous work showing that biodiversity– disease relationships depend on ecological context and species traits [3, 19].

Although it is unsurprising that pathogen transmissibility emerged as the strongest predictor of epidemic intensity, given that transmission potential fundamentally governs pathogen invasion and spread, the more novel result is that timing of K-species establishment strongly influenced pathogen persistence and that its effect depended on pathogen transmissibility. Specifically, delayed establishment of slow life-history hosts had little effect when transmission potential was low but substantially increased persistence for highly transmissible pathogens. These findings suggest that the epidemiological consequences of restoration depend not only on pathogen characteristics, but also on the timing of faunal succession within recovering communities.

### The epidemiological consequences of restoration depend on pathogen transmissibility

The strongest ecological result of this study was the interaction between the timing of slow life-history host establishment and pathogen transmissibility. Delayed establishment of slow lifehistory hosts consistently increased pathogen persistence, but the magnitude of this effect depended strongly on *R*_0_. When pathogen transmissibility was low, persistence remained limited regardless of the timing of slow life-history host establishment. In contrast, highly transmissible pathogens responded strongly to restoration trajectories, with delayed establishment of slow life-history hosts substantially increasing persistence probability. This result provides a potential explanation for why biodiversity–disease relationships often appear inconsistent across ecosystems [3, 19]. Rather than depending solely on community composition, epidemiological outcomes also depend on pathogen transmission potential. Consequently, identical restoration interventions may produce markedly different outcomes depending on pathogen characteristics, consistent with growing evidence that biodiversity effects on disease transmission are contingent on ecological context and species traits [26, 3, 19]. Our results suggest that faunal recovery as a consequence of restoration acts primarily by shortening the period during which competent fast life-history hosts dominate recovering communities [27, 28]. Early establishment of slow life-history hosts reduced pathogen persistence by accelerating this transition, whereas delayed establishment prolonged opportunities for pathogen maintenance. The establishment of slow life-history hosts therefore appears to be an underappreciated lever for disease management, particularly for highly transmissible pathogens. In addition, restoration outcomes are likely to depend on when recovering ecosystems are assessed, as the epidemiological consequences of restoration vary through time with changes in host-community composition. Communities with similar species richness may therefore exhibit different disease outcomes depending on the stage of succession and the functional traits of the species that have established.

### Different mechanisms govern epidemic intensity, persistence, and spread

The relative importance of predictors differed substantially among epidemiological outcomes. Peak prevalence was almost entirely explained by pathogen transmissibility [29], indicating that epidemic intensity is largely determined by transmission potential. In contrast, pathogen persistence was strongly dependent on pathogen transmissibility and timing of community composition dynamics [30]. The timing of epidemic spread followed a different pattern. Transmission between competent fast life-history host populations was one of the strongest predictors of time to epidemic peak. Higher transmission among reservoir populations was associated with later community-wide epidemic peaks, suggesting that infection spread progressively through multiple interconnected host populations rather than producing a rapid outbreak concentrated within a single population. Similar effects have been reported in previous multi-host epidemic models, where host-community structure and transmission pathways strongly influenced epidemic dynamics [31, 32]. Together, these results indicate that epidemic intensity, persistence, and propagation are governed by distinct combinations of epidemiological and ecological processes.

### Community composition matters more than richness alone

Although the timing of species reassembly influenced disease outcomes, maximum species richness consistently exhibited relatively weak predictive importance. Instead, disease dynamics were more strongly associated with the composition and functional characteristics of host communities [33, 34]. In our simulations, restoration modified disease outcomes primarily through the replacement of highly competent fast life-history hosts by less competent slow life-history hosts rather than through increases in species richness alone. This distinction is important because restoration success is often evaluated using richness-based targets [35]. Our results suggest that disease regulation depends more strongly on which host types establish and when they establish [36, 37, 38] than on the total number of species ultimately present. Biodiversity therefore influenced disease dynamics through community assembly processes and changes in host functional structure rather than through richness per se, consistent with growing evidence that biodiversity effects on disease risk emerge through ecological interactions rather than species richness alone [39, 19, 26, 40].

### Implications for restoration planning and One Health

Disease regulation is increasingly recognized as an ecosystem service, yet restoration planning rarely incorporates explicit epidemiological objectives [38, 3]. Our findings suggest that restoration outcomes should not be evaluated solely through biodiversity targets, but also through their effects on pathogen persistence and host-community composition. Several practical implications emerge. First, restoration strategies that accelerate the establishment of slow life-history hosts may reduce long-term pathogen maintenance, particularly for highly transmissible pathogens. Second, disease surveillance may be most effective during early restoration stages when competent fast life-history hosts dominate recovering communities. Third, restoration planning should consider both host-community composition and transmission pathways among competent hosts, as these strongly influence disease dynamics. Notably, maximum species richness had comparatively little predictive power in our analyses, whereas the life-history composition of host communities consistently emerged as a key determinant of disease outcomes. This finding supports a growing shift in restoration ecology from species richness alone towards functional traits and functional diversity as indicators of ecosystem condition and functioning [41]. More broadly, our findings support the integration of disease ecology into ecosystem restoration and One Health frameworks [42, 43]. Successful restoration may depend not only on how much biodiversity is recovered, but also on the pathogen of interest and on how rapidly ecological succession promotes the establishment of functionally distinct slow life-history species that reduce opportunities for pathogen persistence.

### Limitations and future directions

Our framework intentionally simplifies real ecosystems by representing host communities as two functional endpoints of a fast–slow life-history continuum rather than as discrete ecological classes [44, 23]. Consequently, the fast and slow life-history categories used here should be interpreted as abstractions of demographic pace and successional role rather than as species-level classifications. Additional simplifications include the representation of a single pathogen, the absence of pathogen evolution, waning immunity, and repeated external introductions, as well as the use of an implicitly spatial landscape. While these assumptions limit direct extrapolation to particular host–pathogen systems, they allow us to isolate the mechanisms linking restoration dynamics, biodiversity recovery, and disease transmission. Despite these simplifications, the convergence of sensitivity analyses, machine-learning approaches, generalized additive models, bootstrap validation, and SHAP analyses suggests that the main mechanisms identified here are robust. Future work could incorporate explicit spatial structure, heterogeneous host networks, multiple pathogen species, and empirical restoration trajectories to evaluate the generality of these mechanisms across real-world ecosystems.

## Conclusion

Our results demonstrate that pathogen transmissibility and the timing of K-species establishment jointly determine disease dynamics during ecosystem recovery. Pathogen transmissibility largely governs epidemic intensity, whereas restoration timing strongly influences long-term pathogen persistence. By accelerating the replacement of competent fast life-history hosts, restoration can substantially reduce pathogen maintenance, particularly for highly transmissible pathogens. These findings highlight restoration timing and host-community functional composition as underappreciated ecological levers for disease regulation and suggest that successful restoration depends not only on biodiversity recovery itself, but also on the temporal dynamics and functional trajectory of community assembly.

## 5 Acknowlegement

This project was supported by funding from the EU Horizon Scheme HORIZON-CL6-2023-BIODIV-01-17 Grant No.101134969 as part of the RestoreID (Restoring Ecosystems to Stop the Threat of Re-Emerging Infectious Disease). M.V was supported by the BCOMING project (Horizon Europe 415 project 101059483) from the European Union and FNRS.

## A Supplementary Methods

### Population-level transition rates

For a population *i* belonging to strategy *g* ∈ {*r, K*}, the state is (*S_i_, I_i_, R_i_*) and *N_i_* = *S_i_* + *I_i_* + *R_i_*. Births add susceptible individuals at rate *b_g_N_i_*. Deaths occur at rates *d_g_S_i_*, *d_g_I_i_*, and *d_g_R_i_*, and recovery occurs at rate *γ_g_I_i_*. The corresponding transition propensities are summarized below; each event changes only the indicated compartment counts.

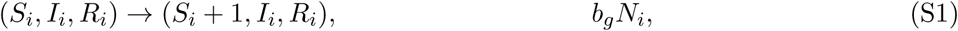

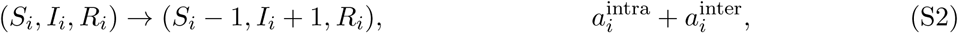

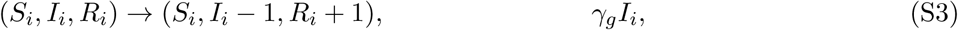

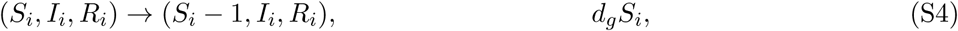

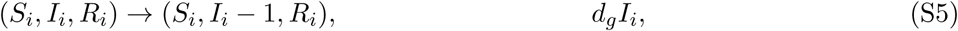

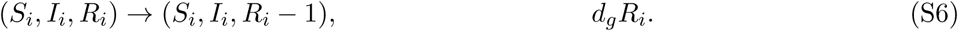

Within-population infection incidence is

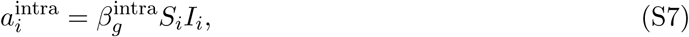

which is density-dependent (mass-action) within each population. Among-population transmission is frequency-dependent with respect to the source group. Let 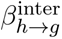 denote the coefficient for transmission from source group *h* ∈ {*r, K*} to recipient group *g*. The implemented coefficients are 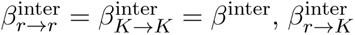 as sampled, and 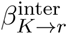 = 0, because no *K*-to-*r* term is implemented. For a recipient population *i* of group *g*, the force of infection is

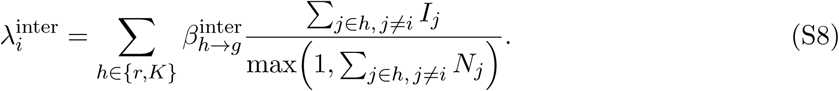

This formulation assumes frequency-dependent transmission among populations, such that transmission depends on the proportion of infectious individuals in the source group rather than its absolute abundance. The exclusion term *j* ≠ *i* prevents a recipient population from contributing to its own inter-population force of infection. The corresponding inter-population infection incidence is given by 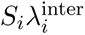. The denominator convention prevents division by zero when a source group is empty. These terms describe community-level transmission potential, not direct human exposure.

### Restoration schedule and strategy assignment

At the beginning of each tau-leap interval, the restoration function returns a target richness *S*_target_(*t*) from the model’s richness routine. The implementation adds populations sequentially until established richness reaches that target; thus, more than one population can arrive at a single update. Arrivals are scheduled by the richness trajectory rather than by an independent stochastic colonization hazard. The richness function is determined by *S*_max_, migration rate, and restoration-time parameters. Before *T_K_*, arrivals are assigned to group *r*. At and after *T_K_*, each arrival is assigned independently to group *K* with probability

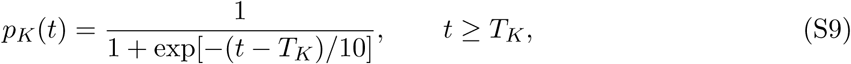

and otherwise to group *r*. Because assignment is evaluated at update times, the timing of arrivals and strategy transitions has resolution *τ*. The first *K* arrival triggers a one-time reduction of each already-established *r* population. The numbers removed are ⌊*S_i_* × *comp*⌋, ⌊*I_i_* × *comp*⌋, and ⌊*R_i_* × *comp*⌋, respectively, where *comp* is a competition parameter representing the proportional reduction of fast life-history host populations following the establishment of slow life-history hosts. The baseline simulations used *comp* = 0.5, whereas robustness analyses explored values ranging from 0.35 to 0.65. Later *r* arrivals are initialized at ⌊*N_i,r_*(1 − *comp*)⌋; earlier *r* populations retain their post-reduction abundances, and *K* arrivals are initialized at *N_i,K_*. These rules are imposed transitions representing the specified succession hypothesis, not emergent competitive dynamics. Effective initial abundances must be positive for all parameter combinations included in analysis. The first population created receives one infectious individual, transferred from susceptible to infectious, when the initialization rule is enabled and the population contains at least one individual.

### Tau-leap algorithm

Simulations use fixed intervals of length *τ* = 0.01, with the final interval shortened if needed to end at *T*_max_. Restoration and species assignments are updated at the beginning of an interval. All demographic and infection propensities are then evaluated from the same pre-leap state, transition counts are sampled, and the changes are applied to obtain the next state. Thus, the implementation is a tau-leap approximation; it does not draw a new global Gillespie waiting time and select one event at a time. For a compartment containing *n* individuals with total exit hazard *h* over an interval of length *τ*, the number of individuals exiting is sampled as

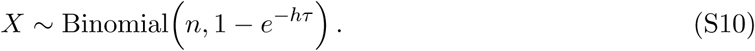

When exits have competing causes with hazards *h*_1_*, . . . , h_q_*, the sampled exits are allocated among causes in proportion to *h_k_/* Σ*_ℓ_ h_ℓ_*. Accordingly, susceptible individuals have infection and death as competing exits, infectious individuals have recovery and death as competing exits, and recovered individuals can die. Births are sampled as

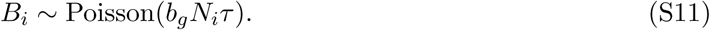

Counts are drawn separately for each population using the pre-leap state, then applied through the corresponding compartment-transition methods. This joint sampling of competing exits prevents the number removed from a compartment from exceeding its pre-leap abundance. Propensities are held fixed within each interval, so accuracy depends on the step size and the rate of state change.

### Initial conditions and epidemiological outcomes

Initial population numbers and abundances are specified by the reference parameter file and the initialization routine. When infection is initialized, one susceptible individual is moved to the infectious compartment so that compartment totals are preserved. Runs with no infection at initialization and no subsequent infection are valid zero-infection realizations; infection extinction does not stop demographic or restoration updates. Metapopulation prevalence is *P* (*t*) = Σ*_i_ I_i_*(*t*)*/* Σ*_i_ N_i_*(*t*) when total abundance is positive, and zero otherwise. For each realization, peak prevalence is defined as the maximum recorded prevalence over [0*, T*_max_] after community richness exceeds two species (*N*_species_ *>* 2), thereby ensuring that epidemic peaks are quantified at the metacommunity level rather than during the initial infection phase of a single host population. Time to peak is the first recorded time at which this maximum is reached, and final prevalence is *P* (*T*_max_). Let *E* = 1 if at least one infectious individual is observed. With numerical tolerance *ϵ*, persistence is coded *Q* = 1 when *E* = 1 and *P* (*T*_max_) *> ϵ*; extinction after infection is coded *D* = 1 when *E* = 1 and *P* (*T*_max_) ≤ *ϵ*. If *E* = 0, both indicators are zero. The tolerance and observation schedule should be held constant across runs.

### LHS design and predictive sensitivity analysis

The LHS design varies the parameters stated for the relevant analysis (see Table. S1): within-population transmission in the *r* and *K* groups, among-population transmission, *K*-group recovery, migration rate, and *T_K_*. In the current sampling configuration, the first five parameters are sampled over 0.5–1.5 times their reference values and *T_K_* over 1–100 model-time units. Any additional parameters or alternative bounds used in a production analysis must be listed explicitly. Integer-valued *T_K_* is sampled on the continuous LHS scale and rounded to an admissible integer before simulation; the realized value is retained in the output. For *n* parameter sets and *R* independent stochastic replicates per set, the design contains *nR* runs. The five-set, one-replicate configuration is a code-testing setup, not a production design for inferential claims. Each run should retain the sampled parameter values, parameter-set and replicate identifiers, seed, *τ*, and outcome values. Seeds should be generated deterministically from the parameter-set and replicate identifiers and passed to the random-number generator used by the simulation. When predictive ranking is performed, continuous outcomes are analyzed with regression and binary persistence/extinction outcomes with classification. Training and test partitions are made by parameter set, keeping all replicates from a parameter combination in the same partition. XGBoost gain and SHAP values describe predictive importance conditional on the sampled parameter space; they are not causal effects.

### Validation and reproducibility

Validation checks include non-negative compartment counts, *N_i_* = *S_i_* + *I_i_* + *R_i_*, consistency of population and metapopulation totals, and prevalence within [0, 1]. Outcome indicators must agree with infection history and final prevalence. Runs without infection remain in the analysis with zero infection outcomes. Tau-leap convergence was evaluated by comparing results across decreasing step sizes. For reproducibility, archive the final source code, parameter file, compiler standard, sampling design and seed, replicate-seed rule, selected *τ*, and output table. Report the actual number of parameter sets and replicates, rather than the testing configuration. The final description of the richness function, parameter bounds, analysis workflow, and numerical checks must match the production code and archived outputs. For reproducibility, the complete source code, parameter files, simulation outputs, and analysis workflow have been archived on Zenodo (DOI: 10.5281/zenodo.23983143). The archived repository includes the compiler standard, Latin hypercube sampling design, parameter ranges, random-seed generation procedure, tau-leaping step size (*τ* = 0.01), and output tables used in the analyses. The descriptions of the richness function, restoration assumptions, parameter bounds, and numerical validation checks reported here correspond exactly to the archived production code and outputs.

## Results

### Illustration of restoration trajectories and stochastic disease dynamics

To illustrate the ecological and epidemiological processes represented in the model, we compared two restoration scenarios differing in the timing of slow life-history host establishment (*T_K_*; Fig. S1). In both cases, total species richness increased progressively throughout the simulation, but the composition of the community differed substantially. When slow life-history hosts were introduced early (*T_K_* = 10; Fig. S1A), their richness increased steadily during community assembly and represented a substantial fraction of the final community. In contrast, when establishment was delayed (*T_K_* = 60; Fig. S1B), communities remained dominated by fast life-history hosts for most of the restoration trajectory and contained relatively few slow life-history hosts by the end of the simulation. Representative stochastic epidemic trajectories further illustrated how restoration timing modified disease outcomes (Fig. S1D). Even under identical parameter values, some realizations resulted in pathogen extinction whereas others persisted throughout the simulation. Early introduction of slow life-history hosts (*T_K_* = 10) was associated with a greater frequency of epidemic fadeout and generally lower long-term prevalence (Fig. S1C). In contrast, delayed introduction of slow life-history hosts (*T_K_* = 60) increased the probability of persistence and produced higher long-term prevalence trajectories (Fig. S1D). These examples illustrate the mechanism identified by the sensitivity analyses: restoration timing modifies disease dynamics by altering the duration of dominance of competent fast life-history hosts within the recovering community.

### Parameter independence within the LHS design

To verify that the Latin hypercube sampling (LHS) procedure did not generate unintended dependencies among predictors, we quantified pairwise Pearson correlations among all sampled parameters (Fig. S2). Correlations were uniformly weak, with all absolute correlation coefficients below 0.07. In particular, epidemiological parameters (*R*_0_, within-species transmission, recovery rate, and among-*r*-species transmission) were effectively independent of restoration-related parameters (migration rate, maximum species richness, and restoration timing). These results indicate that the LHS procedure generated an approximately orthogonal sampling design and minimized collinearity among predictors. The low level of correlation among sampled parameters supports the interpretation of XGBoost variable-importance rankings, as predictor effects are unlikely to be driven by dependencies introduced during parameter sampling.

### Predictive performance and robustness of machine-learning models

The XGBoost models exhibited strong predictive performance across all epidemiological outcomes (Table S2). Predictive accuracy was particularly high for final prevalence and peak prevalence (*R*^2^ *>* 0.92), while time to epidemic peak remained well predicted (*R*^2^ = 0.85). Model performance for pathogen persistence was also high (*R*^2^ = 0.90), indicating that a limited set of ecological and epidemiological parameters captured most of the variation across the simulated restoration scenarios. Bootstrap resampling demonstrated that variable-importance rankings were highly stable (Table S3). Pathogen transmissibility (*R*_0_) consistently emerged as the dominant predictor across all epidemiological outcomes, explaining 73.0% of model gain for final prevalence, 95.4% for peak prevalence, 45.4% for time to peak, and 71.9% for pathogen persistence. Restoration timing (*T_K_*) was the second most important predictor of persistence (23.7%), whereas among-*r* -species transmission was a major determinant of final prevalence (14.2%) and time to peak (25.0%). In contrast, maximum species richness and parameters associated with slow life-history hosts explained comparatively little variation across all outcomes. The strong agreement between bootstrap estimates, SHAP analyses, and gain-based XGBoost rankings indicates that disease dynamics were primarily governed by pathogen transmissibility, while restoration timing and among-*r* -species transmission acted as the main ecological modifiers of persistence and spread.

### SHAP analysis of restoration timing effects on pathogen persistence

SHAP dependence analyses confirmed the strong influence of restoration timing on pathogen persistence identified by the XGBoost models (Fig. S4). The contribution of restoration timing (*T_K_*) to persistence probability increased monotonically across the explored range of values. Early establishment of slow life-history hosts was associated with negative SHAP values, indicating a reduction in pathogen persistence relative to the mean prediction, whereas delayed establishment was associated with increasingly positive SHAP values. The magnitude of this effect depended on pathogen transmissibility. At low values of *R*_0_, restoration timing had relatively limited influence on persistence, and SHAP values remained close to zero across much of the restoration gradient. In contrast, as *R*_0_ increased, delayed introduction of slow life-history hosts produced increasingly positive contributions to pathogen persistence. The largest positive SHAP values were observed when restoration was delayed and pathogen transmissibility was high. These results are consistent with the GAM analysis presented in the main text. Early establishment of slow life-history hosts shortens the period during which highly competent fast life-history hosts dominate the community, thereby reducing opportunities for long-term pathogen maintenance. Conversely, delayed establishment prolongs the dominance of competent hosts and increases persistence probability, particularly for pathogens with high transmission potential.

### Effects of restoration after controlling for pathogen transmissibility

To separate ecological effects from variation in pathogen transmissibility, we repeated the XGBoost analyses under a fixed-*R*_0_ scenario (*R*_0_ = 5; Figure S3). Under the full parameter design, pathogen transmissibility dominated all epidemiological outcomes. In contrast, when *R*_0_ was held constant, restoration timing became the primary predictor of pathogen persistence (55.6% of model gain), whereas its contribution under the unconstrained design was substantially lower (25.1%). The importance of among-*r* -species transmission also increased for several epidemiological outcomes. These results indicate that restoration effects are not solely a consequence of variation in pathogen transmissibility but remain important even when pathogen invasion potential is held constant.

### Robustness to assumptions about competitive replacement

To evaluate whether the inferred drivers of disease dynamics depended on the assumed strength of competitive replacement between fast and slow life-history hosts, we repeated the global sensitivity analysis while varying the competition parameter between 0.35 and 0.65. This parameter determines the proportional reduction of fast life-history host populations following the first establishment of slow life-history hosts. Allowing competition strength to vary did not qualitatively alter model outcomes or predictor rankings (Fig. S5). Pathogen transmissibility (*R*_0_) remained the dominant predictor of final prevalence (42.3%), peak prevalence (60.0%), time to peak (35.8%), and persistence (56.2%). Restoration timing continued to be one of the strongest ecological drivers, particularly for pathogen persistence (23.2%), while among-*r* -species transmission remained important for the timing of epidemic spread (18.6%). The competition parameter itself contributed modestly to model predictions (4.2–8.4% of model gain across outcomes) and did not surpass the influence of pathogen transmissibility, restoration timing, or transmission among competent host populations. These results indicate that the principal conclusions of the study are robust to uncertainty in the magnitude of competitive replacement during succession. In particular, the strong effects of restoration timing and pathogen transmissibility persisted even when the intensity of competition between host functional groups was allowed to vary.

### Does restoration timing act through final community structure?

To determine whether the effect of restoration timing could be explained solely by differences in final community abundance or composition, we fitted GAMs including restoration timing (*T_K_*) together with final abundance (*N*), fast-host richness, or slow-host richness (Table S4). For pathogen persistence, restoration timing remained highly significant in all models (*p <* 0.001), whereas final abundance, fast-host richness, and slow-host richness were not significant predictors. This indicates that the effect of restoration timing on persistence cannot be explained solely by variation in final community abundance or composition. In contrast, both final abundance and host richness significantly contributed to variation in time to epidemic peak. These results suggest that pathogen persistence depends primarily on the trajectory of community assembly, whereas epidemic timing is influenced by both restoration history and the eventual structure of the host community.

---

**Table S1:**
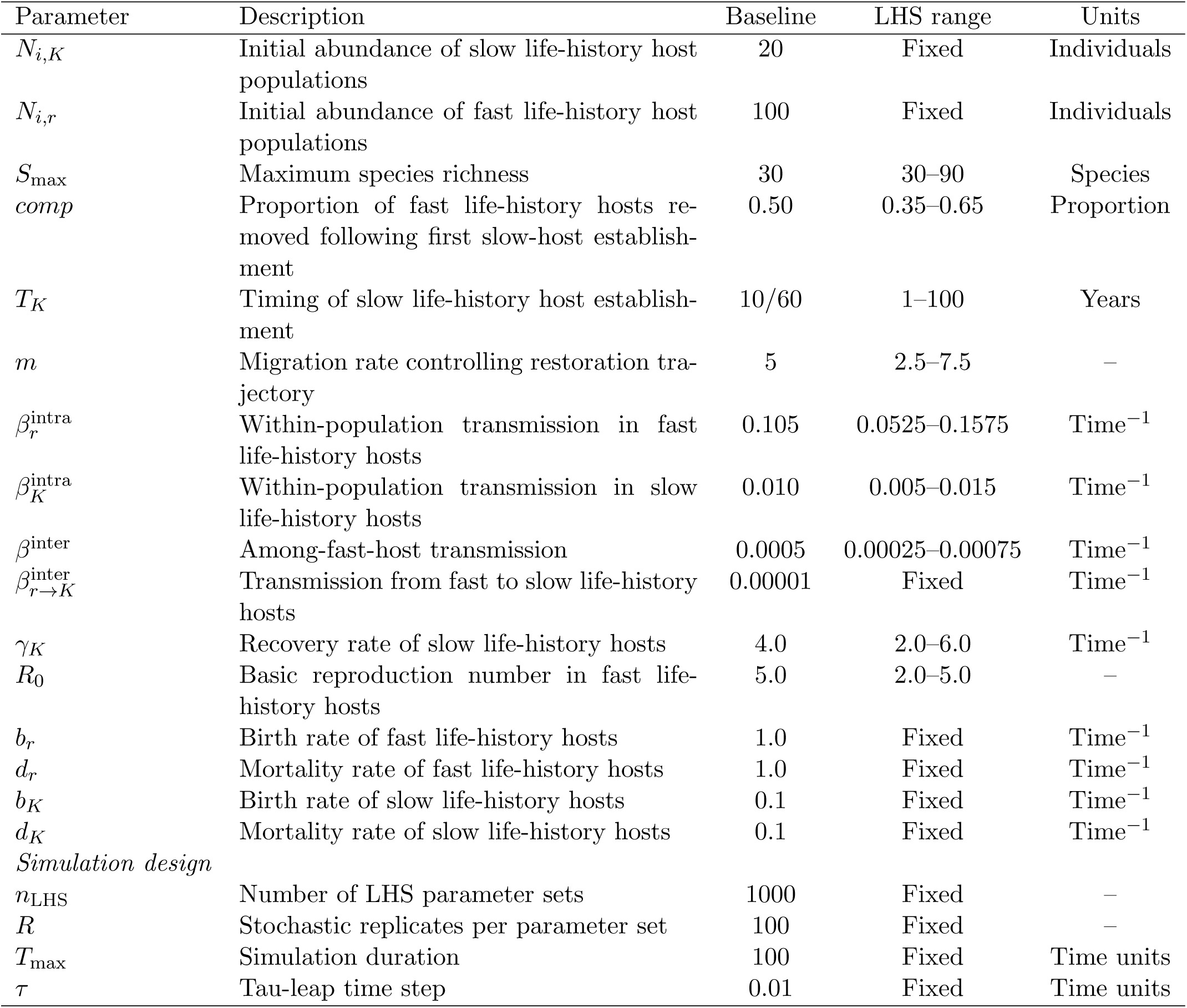
Model parameters and ranges explored in the global sensitivity analysis.

| Parameter | Description | Baseline | LHS range | Units |
|---|---|---|---|---|
| *N*<sub>*i,K*</sub> | Initial abundance of slow life-history host populations | 20 | Fixed | Individuals |
| *N*<sub>*i,r*</sub> | Initial abundance of fast life-history host populations | 100 | Fixed | Individuals |
| *S*<sub>max</sub> | Maximum species richness | 30 | 30–90 | Species |
| *comp* | Proportion of fast life-history hosts re-moved following first slow-host establish-ment | 0.50 | 0.35–0.65 | Proportion |
| *T*<sub>*K*</sub> | Timing of slow life-history host establish-ment | 10/60 | 1–100 | Years |
| *m* | Migration rate controlling restoration tra-jectory | 5 | 2.5–7.5 | – |
| *β*<sub>*r*</sub><sup>intra</sup> | Within-population transmission in fast life-history hosts | 0.105 | 0.0525–0.1575 | Time<sup>−1</sup> |
| *β*<sub>*K*</sub><sup>intra</sup> | Within-population transmission in slow life-history hosts | 0.010 | 0.005–0.015 | Time<sup>−1</sup> |
| *β*<sup>inter</sup> | Among-fast-host transmission | 0.0005 | 0.00025–0.00075 | Time<sup>−1</sup> |
| *β*<sub>*r*→*K*</sub><sup>inter</sup> | Transmission from fast to slow life-history hosts | 0.00001 | Fixed | Time<sup>−1</sup> |
| *γ*<sub>*K*</sub> | Recovery rate of slow life-history hosts | 4.0 | 2.0–6.0 | Time<sup>−1</sup> |
| *R*<sub>0</sub> | Basic reproduction number in fast life-history hosts | 5.0 | 2.0–5.0 | – |
| *b*<sub>*r*</sub> | Birth rate of fast life-history hosts | 1.0 | Fixed | Time<sup>−1</sup> |
| *d*<sub>*r*</sub> | Mortality rate of fast life-history hosts | 1.0 | Fixed | Time<sup>−1</sup> |
| *b*<sub>*K*</sub> | Birth rate of slow life-history hosts | 0.1 | Fixed | Time<sup>−1</sup> |
| *d*<sub>*K*</sub> | Mortality rate of slow life-history hosts | 0.1 | Fixed | Time<sup>−1</sup> |
| *Simulation design* |  |  |  |  |
| *n*<sub>LHS</sub> | Number of LHS parameter sets | 1000 | Fixed | – |
| *R* | Stochastic replicates per parameter set | 100 | Fixed | – |
| *T*<sub>max</sub> | Simulation duration | 100 | Fixed | Time units |
| *τ* | Tau-leap time step | 0.01 | Fixed | Time units |

**Table S2:** Performance of XGBoost models for epidemiological outcomes.

| Outcome | *R*<sup>2</sup> | RMSE |
|---|---|---|
| Final prevalence | 0.928 | 0.0079 |
| Peak prevalence | 0.960 | 0.0113 |
| Time to peak | 0.848 | 2.573 |
| Persistence probability | 0.902 | 0.0518 |

**Figure S1:**
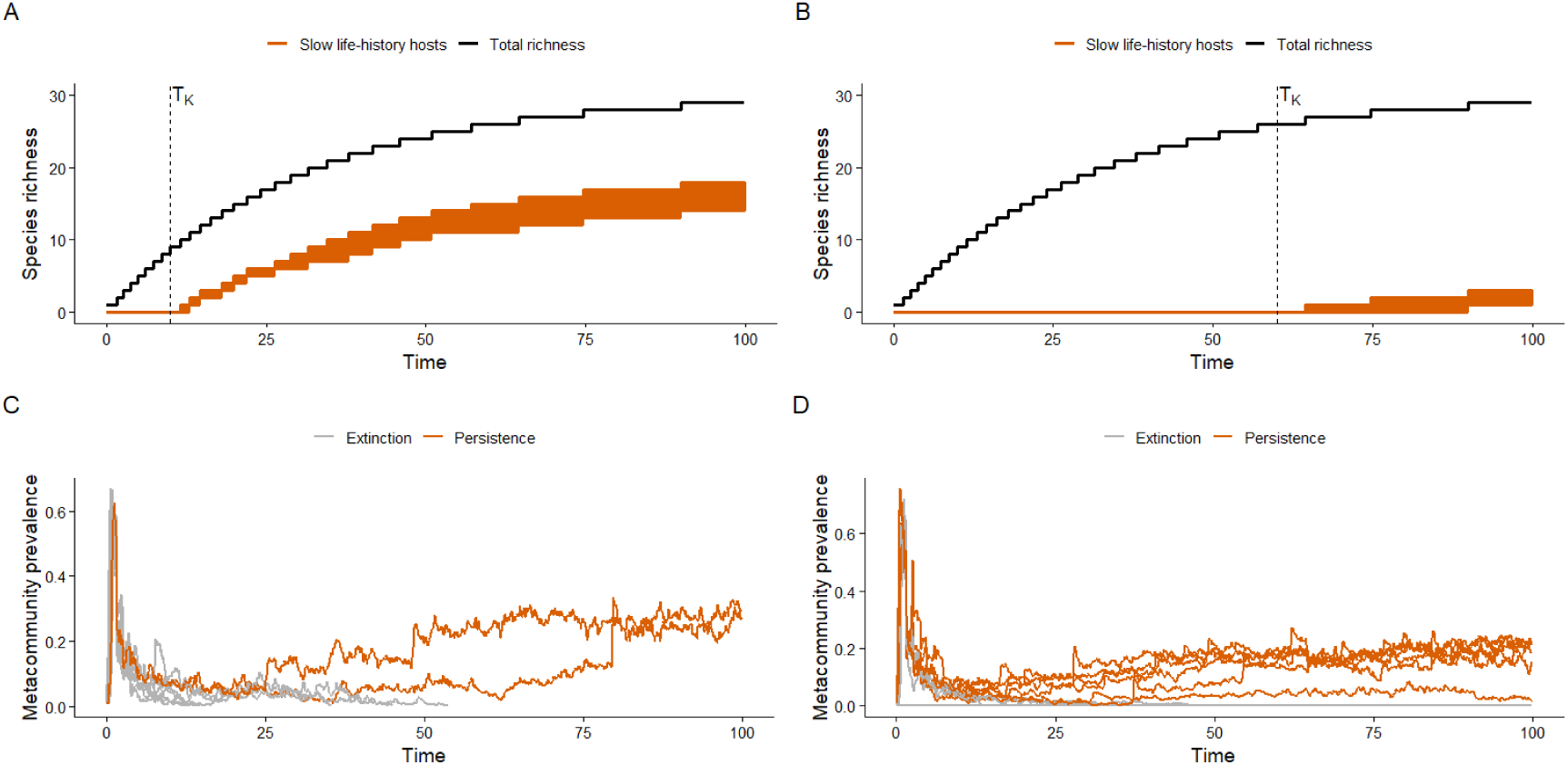
Illustration of restoration trajectories and stochastic epidemic dynamics under contrasting restoration timing scenarios. (A) Community assembly when slow life-history hosts are introduced early (*T_K_* = 10). Black lines show total species richness and orange lines show richness of slow life-history hosts. (B) Community assembly when slow life-history host introduction is delayed (*T_K_* = 60). Dashed vertical lines indicate the restoration threshold *T_K_*. (C) Representative prevalence trajectories from ten stochastic simulations under early introduction of slow life-history hosts. (D) Representative prevalence trajectories from ten stochastic simulations under delayed introduction of slow life-history hosts. Orange trajectories correspond to simulations in which the pathogen persisted until the end of the simulation, whereas grey trajectories correspond to simulations in which the pathogen became extinct. All simulations shown were generated using the same parameter set, including a fixed pathogen transmissibility (*R*_0_ = 5). The only difference between panels A/C and B/D is the timing of slow life-history host introduction (*T_K_* = 10 versus *T_K_* = 60).

**Table S3:** Bootstrap estimates of XGBoost variable importance for each epidemiological outcome. Values correspond to the mean gain across bootstrap replicates, together with standard deviations (SD) and 95% confidence intervals.

| Outcome | Variable | Mean Gain | SD | Lower 95% | Upper 95% |
|---|---|---|---|---|---|
| Final prevalence | *R*<sub>0</sub> of fast life-history hosts | 0.730 | 0.019 | 0.695 | 0.763 |
|  | Among-*r*-species transmission | 0.142 | 0.014 | 0.113 | 0.167 |
|  | Restoration timing (*T*<sub>*K*</sub>) | 0.091 | 0.013 | 0.071 | 0.118 |
|  | Maximum species richness | 0.022 | 0.004 | 0.015 | 0.032 |
| Peak prevalence | *R*<sub>0</sub> of fast life-history hosts | 0.954 | 0.003 | 0.947 | 0.961 |
|  | Migration rate | 0.027 | 0.003 | 0.022 | 0.033 |
|  | Among-*r*-species transmission | 0.006 | 0.001 | 0.005 | 0.007 |
|  | Restoration timing (*T*<sub>*K*</sub>) | 0.003 | 0.001 | 0.002 | 0.004 |
| Time to peak | *R*<sub>0</sub> of fast life-history hosts | 0.454 | 0.026 | 0.410 | 0.510 |
|  | Among-*r*-species transmission | 0.250 | 0.022 | 0.207 | 0.288 |
|  | Restoration timing (*T*<sub>*K*</sub>) | 0.105 | 0.022 | 0.063 | 0.153 |
|  | Maximum species richness | 0.093 | 0.018 | 0.066 | 0.132 |
| Persistence probability | *R*<sub>0</sub> of fast life-history hosts | 0.719 | 0.017 | 0.689 | 0.750 |
|  | Restoration timing (*T*<sub>*K*</sub>) | 0.237 | 0.015 | 0.204 | 0.267 |
|  | Among-*r*-species transmission | 0.015 | 0.002 | 0.011 | 0.020 |
|  | Migration rate | 0.008 | 0.001 | 0.006 | 0.012 |

**Table S4:**
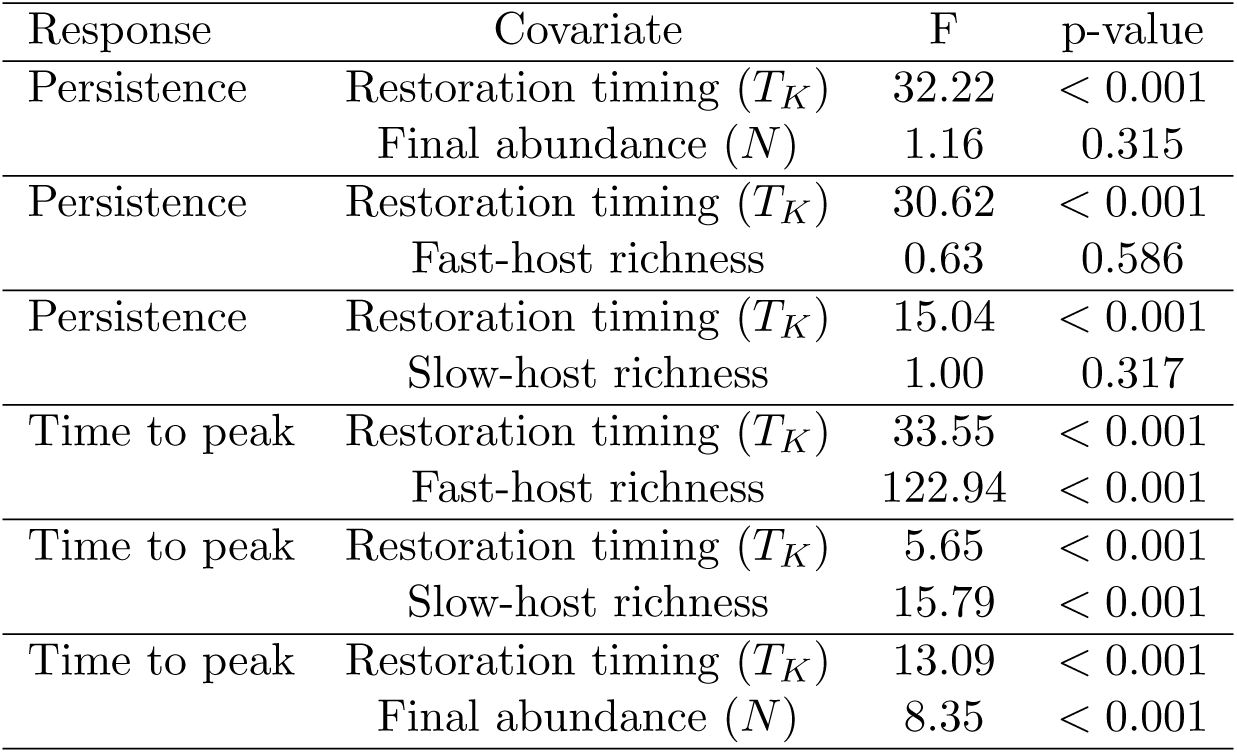
Generalized additive models evaluating whether the effect of restoration timing (*T_K_*) can be explained by final community abundance or composition.

| Response | Covariate | F | p-value |
|---|---|---|---|
| Persistence | Restoration timing (*T<sub>K</sub>*) | 32.22 | < 0.001 |
|  | Final abundance (*N*) | 1.16 | 0.315 |
| Persistence | Restoration timing (*T<sub>K</sub>*) | 30.62 | < 0.001 |
|  | Fast-host richness | 0.63 | 0.586 |
| Persistence | Restoration timing (*T<sub>K</sub>*) | 15.04 | < 0.001 |
|  | Slow-host richness | 1.00 | 0.317 |
| Time to peak | Restoration timing (*T<sub>K</sub>*) | 33.55 | < 0.001 |
|  | Fast-host richness | 122.94 | < 0.001 |
| Time to peak | Restoration timing (*T<sub>K</sub>*) | 5.65 | < 0.001 |
|  | Slow-host richness | 15.79 | < 0.001 |
| Time to peak | Restoration timing (*T<sub>K</sub>*) | 13.09 | < 0.001 |
|  | Final abundance (*N*) | 8.35 | < 0.001 |

**Figure S2:**
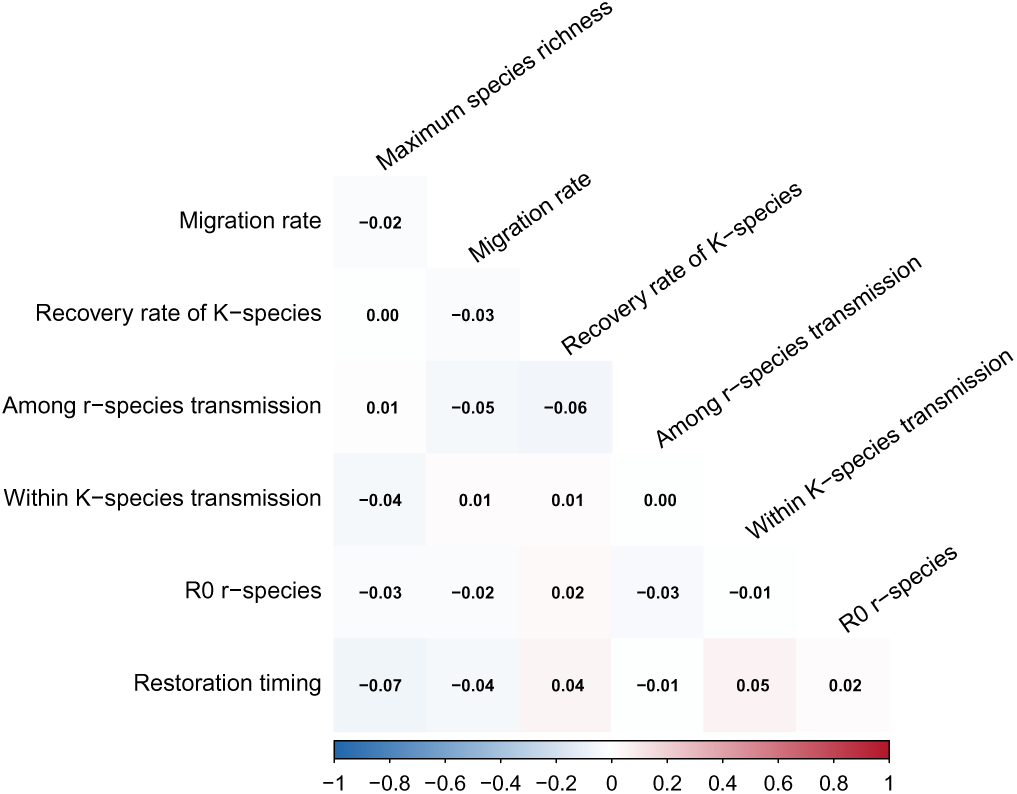
Pairwise Pearson correlations among parameters sampled in the Latin hypercube design.

**Figure S3:**
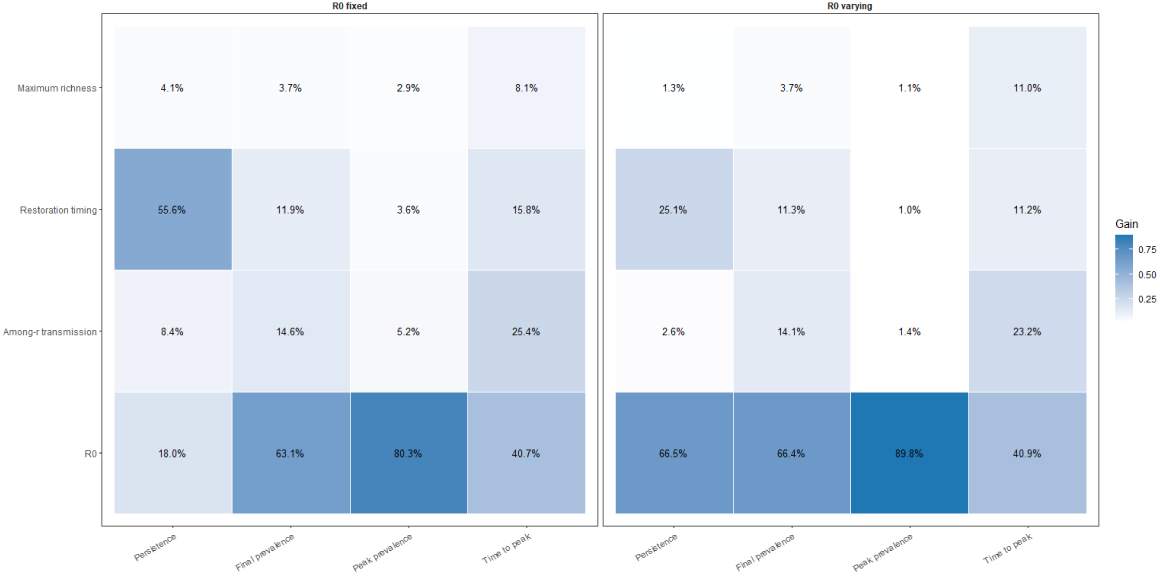
Comparison of XGBoost variable-importance rankings obtained under the full parameter sampling design (variable *R*_0_) and under a constrained transmission scenario (*R*_0_ = 5). Values represent the relative contribution (gain) of each predictor to model performance for persistence probability, final prevalence, peak prevalence, and time to peak. Fixing *R*_0_ reduced the importance of pathogen transmissibility and increased the relative contribution of ecological drivers, particularly restoration timing.

**Figure S4:**
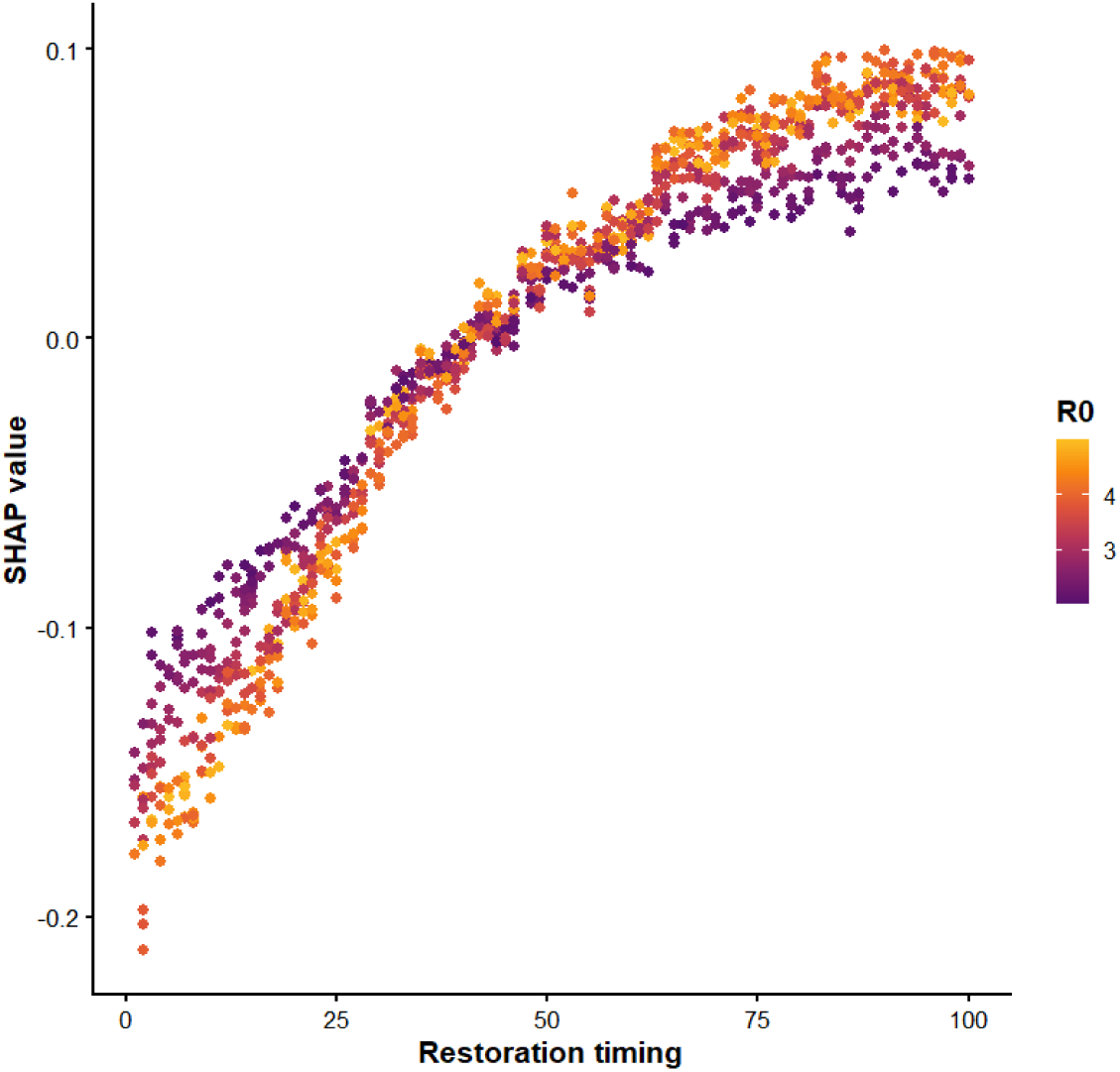
SHAP dependence plot showing the effect of restoration timing (*T_K_*) on pathogen persistence probability. Points represent individual parameter sets and are coloured according to pathogen basic reproduction number (*R*_0_). Positive SHAP values indicate that higher values of restoration timing contribute positively to predicted persistence, whereas negative values indicate a reduction in persistence.

**Figure S5:**
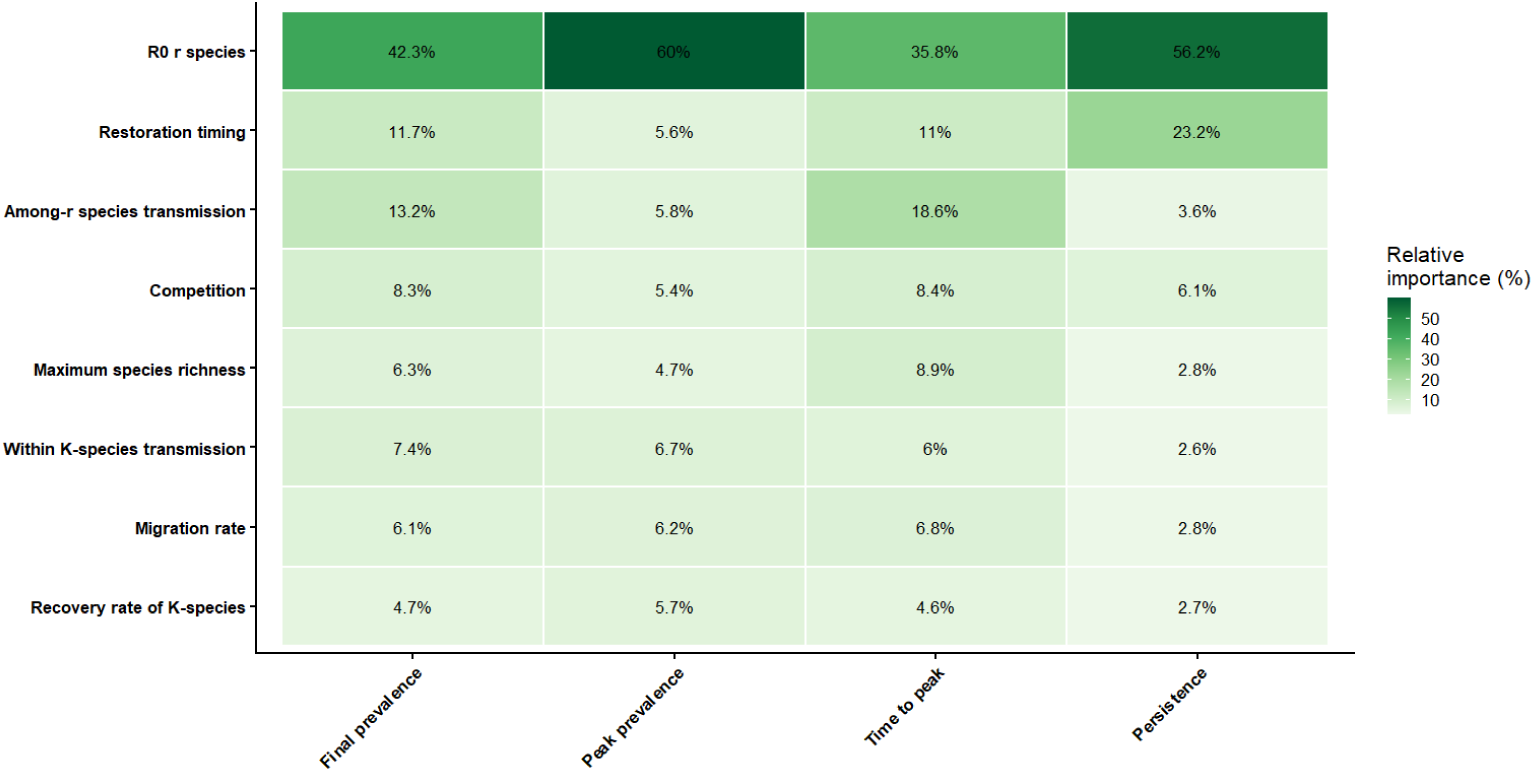
Robustness analysis of XGBoost variable importance rankings when the strength of competitive replacement between fast and slow life-history hosts was varied between 0.35 and 0.65. Cell values indicate the percentage contribution (gain) of each predictor to model performance for final prevalence, peak prevalence, time to peak, and persistence probability. Despite variation in competition strength, pathogen transmissibility (*R*_0_), restoration timing, and among-*r* -species transmission remained the dominant predictors, indicating that the main conclusions of the study are robust to uncertainty in competitive replacement during succession.

